# Stochastic dynamics of consumer-resource interactions

**DOI:** 10.1101/2021.02.01.429174

**Authors:** Abhyudai Singh

## Abstract

The interaction between a consumer (such as, a predator or a parasitoid) and a resource (such as, a prey or a host) forms an integral motif in ecological food webs, and has been modeled since the early 20*^th^* century starting from the seminal work of Lotka and Volterra. While the Lotka-Volterra predator-prey model predicts a neutrally stable equilibrium with oscillating population densities, a density-dependent predator attack rate is known to stabilize the equilibrium. Here, we consider a stochastic formulation of the Lotka-Volterra model where the prey’s reproduction rate is a random process, and the predator’s attack rate depends on both the prey and predator population densities. Analysis shows that increasing the sensitivity of the attack rate to the prey density attenuates the magnitude of stochastic fluctuations in the population densities. In contrast, these fluctuations vary non-monotonically with the sensitivity of the attack rate to the predator density with an optimal level of sensitivity minimizing the magnitude of fluctuations. Interestingly, our systematic study of the predator-prey correlations reveals distinct signatures depending on the form of the density-dependent attack rate. In summary, stochastic dynamics of nonlinear Lotka-Volterra models can be harnessed to infer density-dependent mechanisms regulating consumer-resource interactions. Moreover, these mechanisms can have contrasting consequences on population fluctuations, with predator-dependent attack rates amplifying stochasticity, while prey-dependent attack rates countering to buffer fluctuations.

## I. Introduction

Consumer-resource dynamics has been traditionally studied using an ordinary differential equation framework starting from the seminal work of Lotka and Volterra over a century ago [1]–[7]. The classical Lotka-Volterra model

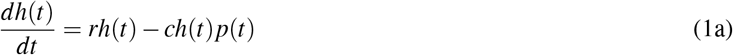

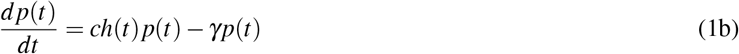

captures the dynamics of a predator-prey system, where *h*(*t*) and *p*(*t*) are the average population densities (number of individuals per unit area) of the prey (i.e., the resource), and the predator (i.e., the consumer), at time *t*. Here *r* represents the prey’s growth rate and *h*(*t*) grows exponentially over time in the absence of the predator. Predators consume prey with rate *c* that we refer to as the *attack rate*, and each attacked prey leads to a new predator. Finally, each predator dies at a rate γ. In addition to predator-prey systems, ecological examples of such consumer-resource dynamics include host-parasitoid interactions that have tremendous application in biological control of pest species [8]–[15]. In a typical interaction, parasitoid wasps search and attack their host insect species by laying an egg within the body of the host. The egg hatches into a juvenile parasitoid that develops within the host by eating it from the inside out. Once fully developed, the parasitoid emerges from the dead host to repeat the life cycle.

The steady-state prey and predator equilibrium densities corresponding to the Lotka-Volterra model (1) are given by

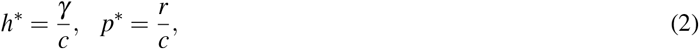

respectively. It turns out that this equilibrium is neutrally stable resulting in cycling population densities with a period of 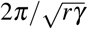 (assuming perturbations around the equilibrium) [16], and such population cycles have fascinated theoretical ecologists with several interpretations/extensions [17]–[19]. There is a rich body of literature expanding the Lotka-Volterra model to understand how diverse processes can push the equilibrium towards stability or instability [20]–[29]. For example, self-limitation in the prey’s growth in the form of a carrying capacity stabilizes the equilibrium [16]. Interestingly, a wide class of two-dimensional consumer-resource models with an unstable equilibrium result in a stable limit cycle [30], [31].

In this contribution, we focus on generalizing (1) to

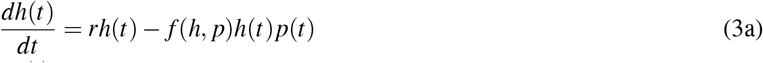

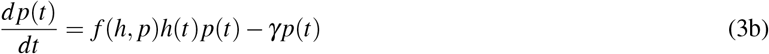

that considers a density-dependent attack rate *f* (*h, p*), where *f* is a continuously differentiable function in both arguments. A generalized attack rate encompasses a wide range of ecological mechanisms. At one end of the spectrum are prey-dependent attack rates that capture nonlinear functional responses. For a Type II functional response

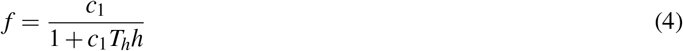

is a decreasing function of the prey density, where *c*_1_ > 0 is the attack rate at small prey densities and *T_h_* is the handling time [32]–[34]. Basically, the total attack rate per predator *f* (*h, p*)*h* increases linearly with *h* at low prey densities, but saturates to 1/*T_h_* at high prey densities. Similarly, a Type III functional response corresponds to a sigmoidal function

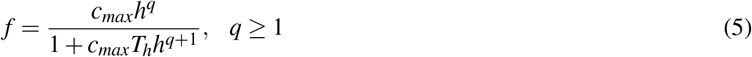

that initially accelerates with increasing prey density and then saturates to 1/*T_h_*. At the other end of the spectrum are predator-dependent attack rates. For example, a decreasing attack rate with increasing predator density implies mutual interference between predators [35]–[38], or aggregation of predators to a subpopulation of high-risk individuals [39]–[43]. In contrast, cooperation between predators is reflected in *f* increasing with predator density.

In this contribution we provide analytical conditions for having a stable population dynamics in terms of the sensitivity of *f* to prey/predator densities, thus combining the impact of different mechanisms into a single generalized stability criterion. Furthermore, we consider a stochastic formulation of the model by allowing the prey’s growth rate to follow a OrnsteinUhlenbeck random process that drives the deterministic predator-prey dynamics (3). We systematically investigate how random fluctuations in the prey’s growth rate propagate to population densities and uncover mechanisms that amplify or attenuate these fluctuations.

## II. Stability analysis of the generalized Lotka-Volterra model

Setting the left-hand-side of (3) to zero, the equilibrium population densities *h*\* and *p*\* of the generalized Lotka-Volterra model are the solution to

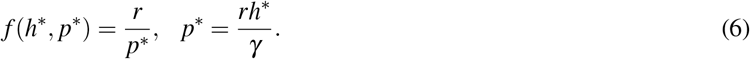

We assume that the function form of *f* is such that (6) yields a *unique* non-trivial equilibrium. Before performing a local stability analysis around the equilibrium, we define two dimensionless log sensitivities 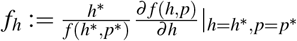, log sensitivity of the attack rate *f* (*h,p*) to the prey density 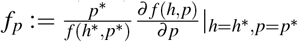, log sensitivity of the attack rate *f*(*h,p*) to the predator density where 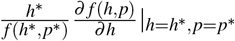 is the partial derivative of *f* with respect to *h* evaluated at the equilibrium. To be biology realistic, we assume that *f*(*h, p*)*p* is an increasing function of the predator density that constrains *f_p_* > −1, i.e., the decrease in *f* with increasing *p* cannot be faster than 1/*p*.

Linearizing the right-hand-side of (3) around the equilibrium yields the following Jacobian matrix

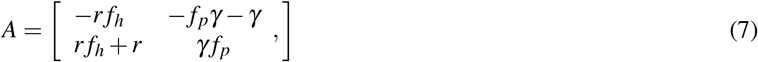

and stability requires a Hurwitz matrix whose eigenvalues have negative real parts [44], [45]. For a two-dimensional system, the equilibrium is asymptotically stable, if and only if, the determinant of the *A* matrix is positive and its trace is negative [44], [45]. This implies that the equilibrium obtained as the solution to (6) is asymptotically stable, if and only if, both these inequalities hold

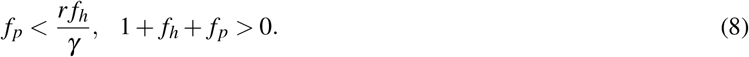

The black shaded region in Fig. 1 shows the stability region as a function of the log sensitivities *f_p_* and *f_h_*, with the neutrally stable Lotka-Volterra equilibrium corresponding to *f_p_* = *f_h_* = 0 on the edge of stability. Moreover, the intersection of the two lines in Fig. 1 reveals

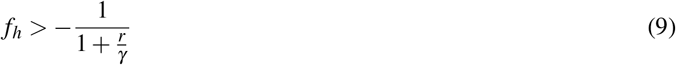

as a necessary condition for stability. These stability conditions for continuous-time consumer-resource models are analogous counterparts to recently developed stability conditions for discrete-time consumer-resource models [46], [47].

**Fig. 1:**
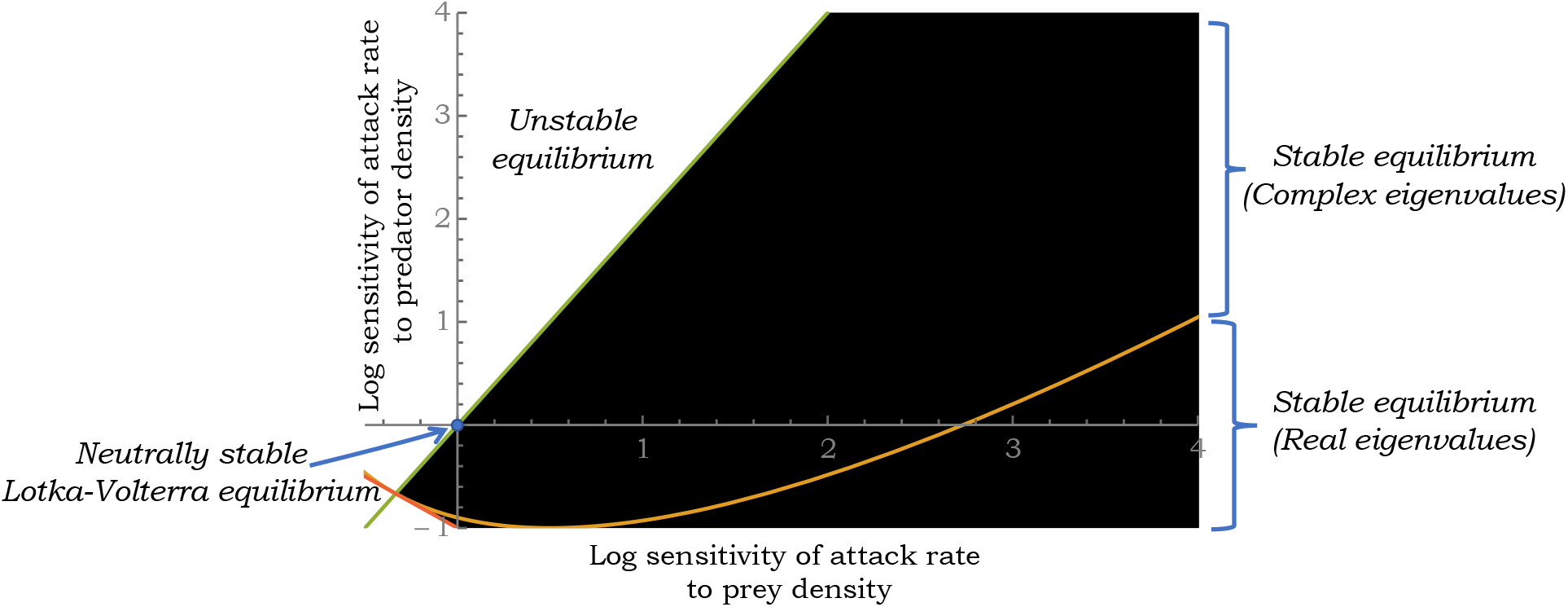
The black shaded region represents the region of stability for the equilibrium of the generalized Lotka-Volterra model (3). The stability criterion (10) is plotted in terms of the log sensitivities of the attack rate *f* (*h, p*) to the prey (*f_h_*) and predator (*f_p_*) densities. The yellow line within the stability region separates the regions of negative real eigenvalue of the Jacobian matrix and complex eigenvalues with negative real parts. For this plot, the prey’s growth rate is assumed to be *r* = 2 per unit time and *γ* = 1 per unit time.

It’s clear from the Fig. 1 that as reported in previous analysis [48], a Type II functional response with *f_h_* < 0 and *f_p_* = 0 will lead to an unstable equilibrium. In contrast, *f_h_* > 0 and *f_p_* = 0 stabilizes the equilibrium. As discussed earlier, *f_h_* > 0 arises in the initial phase of a Type III functional response where the predator attack rate accelerates with increasing prey density. Interestingly, a Type II functional response (*f_h_* < 0) can provide stability in a narrow range if combined with other mechanisms, such as mutual interference between predators where *f_p_* < 0. overall these results show that an attack rate that increases with prey-density is sufficient to stabilize the equilibrium as long as −1 < *f_p_* < *r* * *f_h_*. Similarly, a predator-dependent attack rate with *f_h_* = 0 and −1 < *f_p_* < 0 is sufficient to stabilize the equilibrium. Finally, we point out that the line

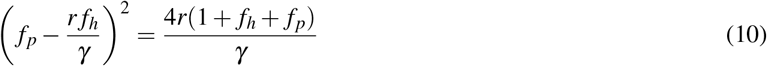

divides the stability region into two parts – negative real eigenvalues of the Jacobian matrix below the line, and complex eigenvalues with negative real parts above the line. With increasing *r*, the line shifts further to the left. This separation within the stability region is relevant in the stochastic formulation of the model, where a stable equilibrium with complex eigenvalues of the *A* matrix can yield signatures of oscillatory dynamics in the presence of noise.

## III. Stochastic formulation of the generalized Lotka-Volterra model

Having determined the stability regions of the deterministic model (3), we next turn our attention to the stochastic formulation of this model. In this context, much prior work has studied demographic stochasticity arising at low population abundances using Lotka-Volterra and spatial predator-prey models [44], [49]–[54]. Here, we focus on environmental stochasticity that arises through randomness in the prey’s growth rate. Towards that end, we let the prey’s growth rate *r*(*t*) evolve as per an Ornstein-Uhlenbeck (OU) process

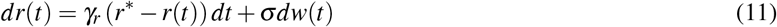

where *w*(*t*) is the Wiener process and *r*\* is the mean level of *r*(*t*). By using an OU process we capture memory in growth-rate fluctuations, with parameters *γ_r_* > 0 and σ > 0 characterizing the time-scale and magnitude of *r*(*t*) fluctuations, respectively. These growth-rate fluctuations in turn drive population-density fluctuations through the model

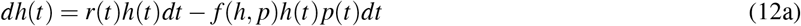

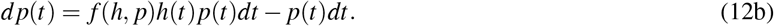

Before describing mathematical tools for quantifying statistical moments of population densities, we point out that our approach of incorporating environmental stochasticity is different to other works that have considered either seasonal deterministic variations in *r*(*t*) [44], [55] or have added memoryless Brownian noise terms to the deterministic population dynamics [56]–[59].

To obtain the time evolution of the statistical moments of *r*(*t*), *h*(*t*) and *p*(*t*) corresponding to the nonlinear stochastic dynamical system (11)-(12) we use the following result. For any continuously differentiable function *ψ*(*r,h,p*), its expected value evolves as

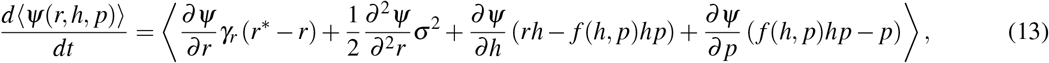

and moment dynamics is obtained by simply using a monomial

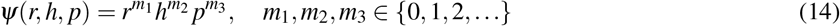

in (13) [60]. Throughout the manuscript we use { } to denote the expected value operation. For example, taking *m*_1_ = 1 or 2 with *m*_2_ = *m*_3_ = 0 yields the time evolution of the first two statistical moments of *r*(*t*)

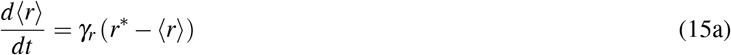

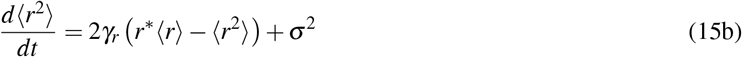

that result in the following steady-state mean and variance

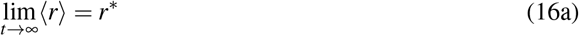

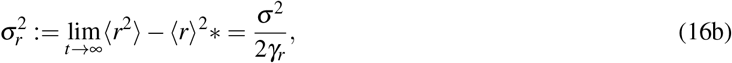

respectively. Similarly, to derive the mean dynamics of the prey’s population density we use *m*_2_ = 1, *m*_1_ = *m*_3_ = 0 to obtain

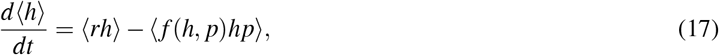

where the right-hand-side now consists of higher-order moments. This problem of unclosed moment dynamics, where the time evolution of lower-order moments depends on higher-order moments has been well described for nonlinear stochastic systems, and often arises in the modeling of biochemical and ecological processes [15], [61]–[77]. Typically, different closure schemes are employed to approximate moment dynamics and we use one such approach known as the Linear Noise Approximation (LNA) [78]–[82]. In essence, assuming a stable equilibrium (*h*\*, *p*\*) in the deterministic formulation as given by (replacing *r* by *r*\* in (6))

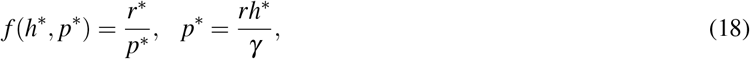

then for small fluctuations in *r*(*t*), *h*(*t*), *p*(*t*) around their respective equilibriums, the model nonlinearities can be linearized as

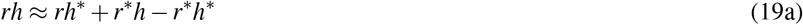

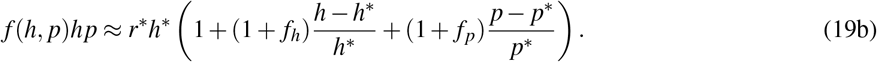

Moment dynamics is then derived after replacing these linear approximations in place of their nonlinear terms in (13) resulting in a closed system – the time derivative of a second-order moment now only depends on moments of order up to two. More specifically, if we collect all the first and second-order moments within the vector

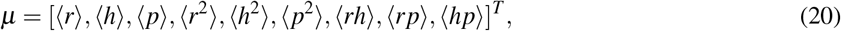

then it’s time evolution is given by a linear time-invariant system

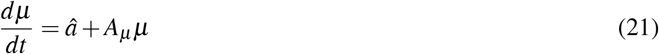

for some *â* and matrix *A_μ_*. Solving this linear system at steady-state quantifies the magnitude of fluctuations in the population densities.

## IV. Results and Discussion

In the previous section, we described a LNA-based approach to quantify the statistical moments of population densities. Here we present some of the key results and insights from the analysis of moments. We quantify noise in the random processes *r*(*t*), *h*(*t*), *p*(*t*) using the square of their respective coefficient of variations

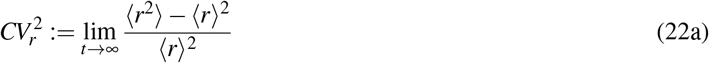

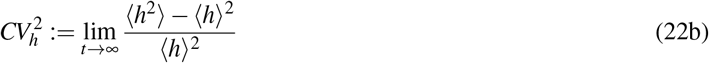

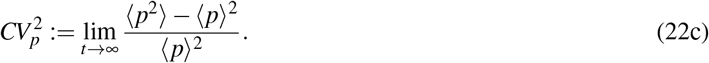

Solving (21) in in Mathematica [83] yields the following analytical expressions for the noise in the prey and the predator population densities (normalized by the noise in the prey’s growth rate)

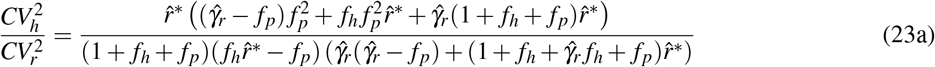

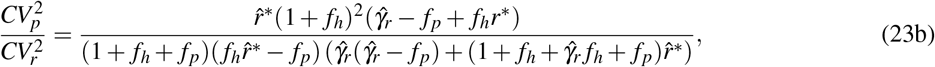

respectively, and they depend on four dimensionless parameters – the sensitivity of the attack rate to the prey density (*f*_h_), the sensitivity of the attack rate to the predator density (*f*_p_), the prey’s average growth rate and the time-scale of fluctuations in *r*(*t*) normalized by the predator’s death rate

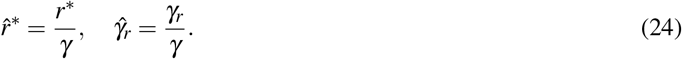

Recall from (10) that the stability of the deterministic equilibrium constrains *f_h_* and *f_p_* in the stability region of Fig. 1 which ensures positivity of noise levels. Moreover, as one gets closer to the Lotka-Volterra model (*f_h_* → 0 and *f_p_* → 0), the system approaches the stability boundary leading to (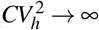 and 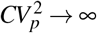) in the LNA framework of noise derivation.

For an attack rate that only depends on the prey’s density (*f_p_* = 0), (23) reduces to

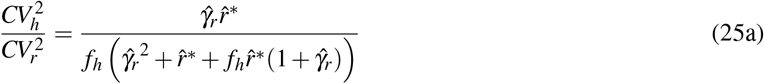

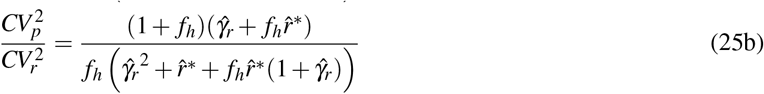

and fluctuations in population densities monotonically decrease with increasing dependence of the attack rate on the prey’s density (Fig. 2). Interestingly, while

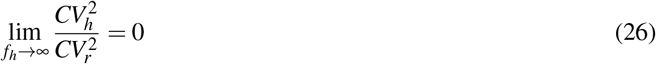

the noise in the predator’s abundance approached a non-zero limit illustrating noise propagation of growth-rate fluctuations to the predator density via the prey.

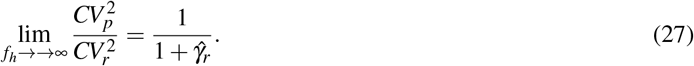

**Fig. 2:**
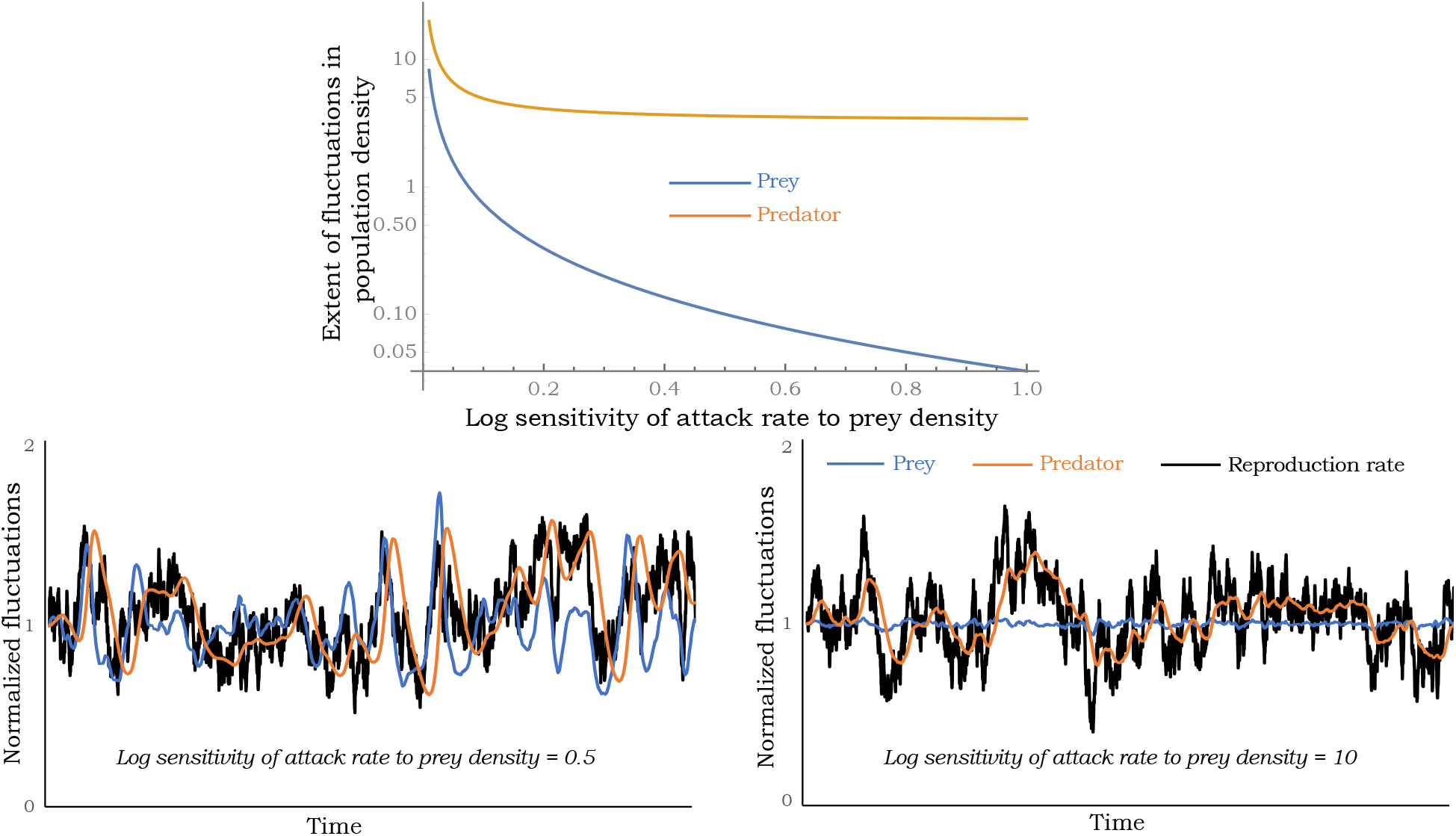
Noise in the fluctuations of population densities as determined by (25) plotted for increasing *f_h_* for *f_p_* = 0. Stochastic realizations of the prey’s growth rate, population densities of the prey and the predator are shown for *f_h_* = 0.5 (left) and *f_h_* = 10 (right). Note that for large values of *f_h_* fluctuations in the predator density remain pronounced even though fluctuations in the prey density are minimal. For this plot, 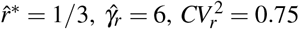

Our analysis further shows that when 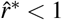, then 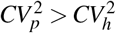. In contrast, when 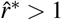, then 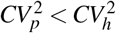 for small value of *f_h_*, and 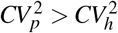 beyond a critical value *f_h_*. One observation from the equilibrium analysis in (18) is that for a prey-dependent attack rate, *h*\* is independent of *r*\* implying that if the prey’s growth rate is chosen from a static distribution then it will not create any fluctuations in the preys’ density. This can be seen in (25) where in the limit

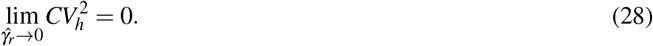

However, as *p*\* is linearly dependent of *r*\*

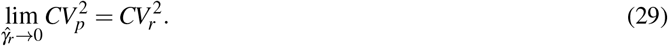

While fluctuations in population densities monotonically decrease with increasing *f_h_*, the impact of a predator-dependent attack rate is quite different with noise levels varying non-monotonically with *f_p_* (Fig. 3). This effect can be understood in term of the stability region in Fig. 1, where for a given *f_h_* > 0 stability requires −1 – *f_h_* < *f_p_* < *r*\* *f_h_*, and increasing *f_p_* on either side puts the system closer to the stability boundary amplifying random fluctuations. This results in a scenario where fluctuations in population densities are minimized at an intermediate value of *f_p_*.

**Fig. 3:**
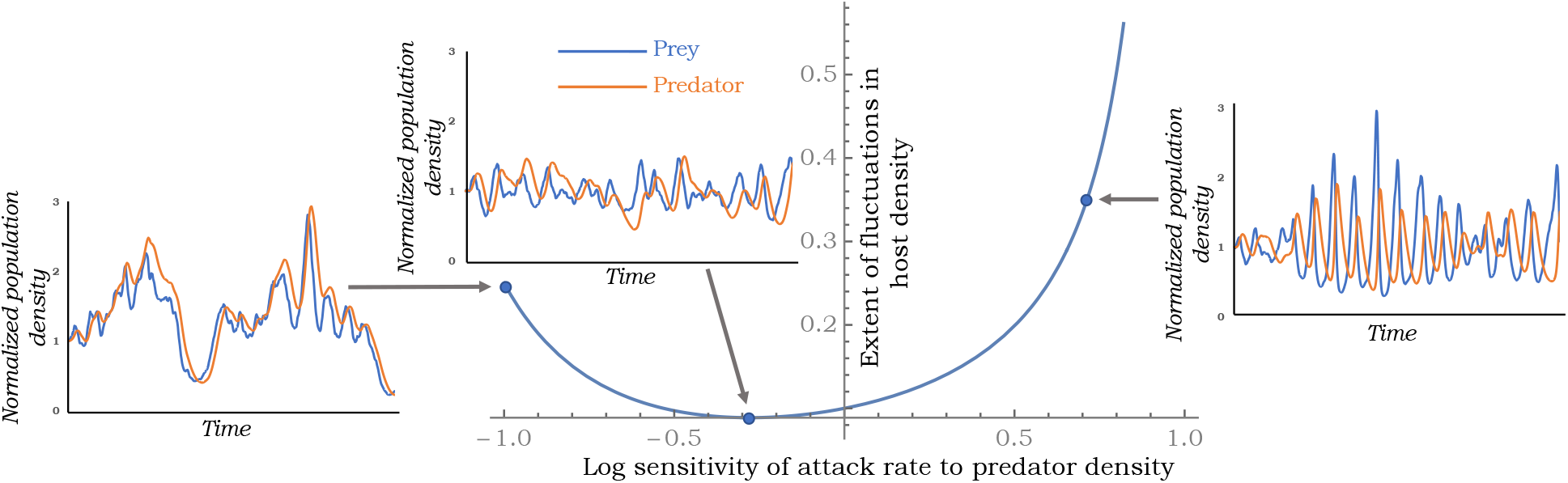
Noise in the fluctuations of prey density 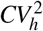 as determined by (23) plotted as a function of *f_p_*. Noise is minimized at an intermediate value of *f_p_* and stochastic realizations of the prey and predator densities are shown for three different values of *f_p_*. For this plot, 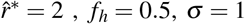 and 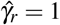.

We next investigate the predator-prey Pearson’s correlation coefficient

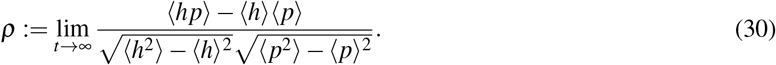

The moment dynamics (21) results in the following closed-form expression

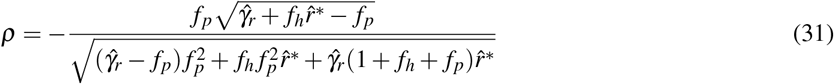

highlighting an interesting result – while a negative dependence of the attack rate (*f_p_* < 0) on the predator density leads to positive predator-prey correlations, a positive dependence *f_p_* > 0 leads to negative correlations (Fig. 4). Moreover, predator-prey density fluctuations are predicted to be uncorrelated for a prey-dependent attack rate. To understand this result, one can derive from (18) the following log sensitivities of the equilibrium densities to *r*\*

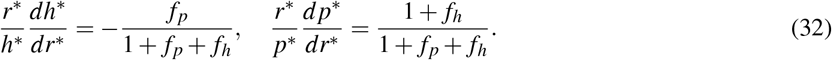

**Fig. 4:**
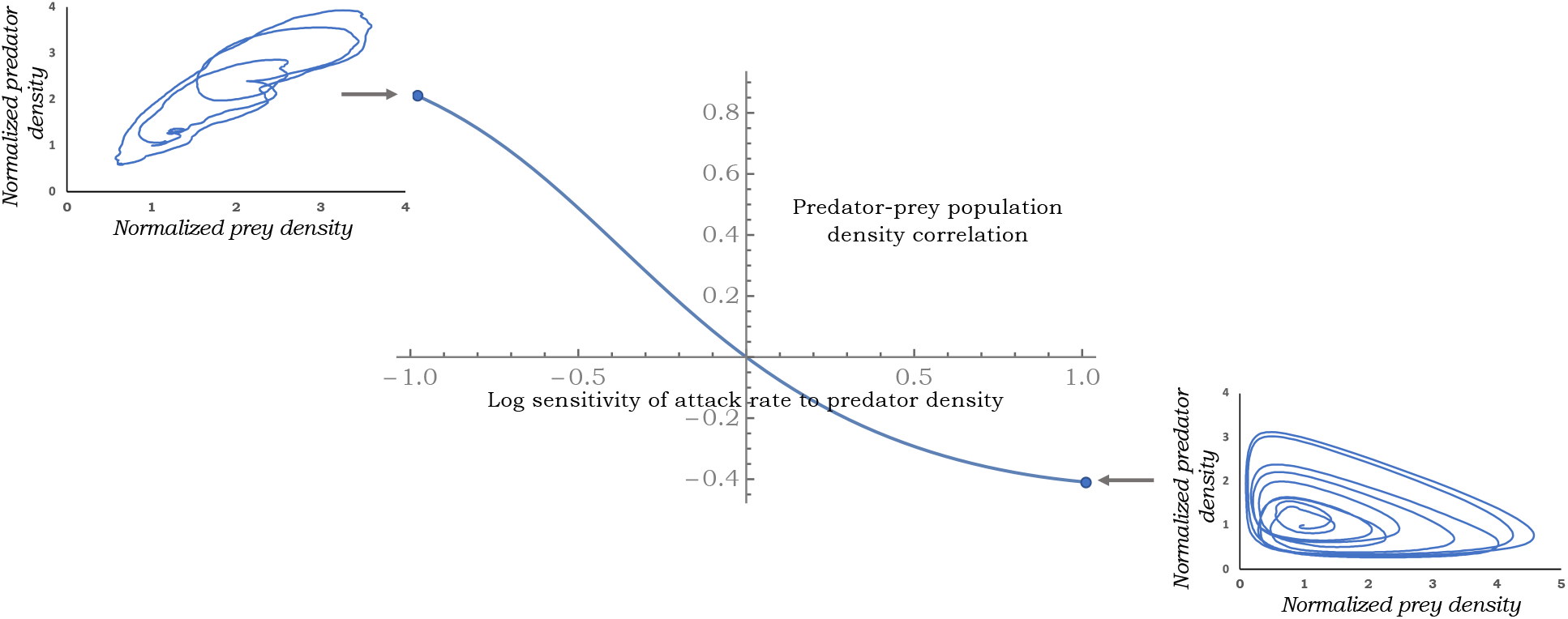
Pearson’s correlation coefficient between the predator and the prey population densities as predicted by (31) for varying levels of *f_p_* with *f_p_* < 0 (*f_p_* > 0) driving a positive (negative) correlation. Sample trajectory paths are shown for *f_p_* = 1 and *f_p_* = −1. Other parameters taken as 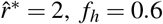 and 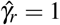.

Thus, when *f_p_* > 0, the prey’s equilibrium density decreases with increasing *r*\*, while the predator’s equilibrium density always increases with *r*\*. These opposing responses of equilibrium densities intuitively explain the negative correlation seen for *f_p_* > 0. In contrast, when *f_p_* < 0, both equilibrium densities increase with *r*\* and manifest in a positive correlation in the stochastic model. Recent work in host-parasitoid discrete-time models with a random host reproduction rate has also identified contrasting correlations depending on the mechanism stabilizing the population dynamics [84].

In summary, we have developed a novel stability criterion for a generalized Lotka-Volterra model with a density-dependent attack rate. (Fig. 1). These result reveal that a Type II functional response can stabilize the equilibrium if combined with mechanisms involving predator inefficiency that puts the system in the black shaded region corresponding to *f_h_* < 0 and *f_p_* > 0. Moreover, stability arises quite robustly for a Type III functional response when *f_h_* > 0 as long as *f_p_* is small enough to be in between −1 – *f_h_* and *rf_h_*. This stability is also reflected in the stochastic model where density fluctuations monotonically decrease with increasing *f_h_* for a prey-dependent attack rate (Fig. 2). However, predator-dependent attack rates can amplify stochasticity as *f_p_* gets close to the stability boundary on either side of the x-axis in Fig. 1. Finally, we have shown that population density correlations may contain signatures on stabilizing mechanisms at play with no correlation in predator-prey densities implying a prey-dependent attack rate (Fig. 4), a negative correlation implying predator cooperation (*f_p_* > 0), and a positive correlation implying mutual interference between predators (*f_p_* < 0). Future work will expand these results to consider demographic stochasticity by explicitly modeling probabilistic birth-death events. It will also be interestingly to consider competition between two or more consumers, and also look at apparent competition between different prey species attacked by a common predator [85], [86]. Along these lines, new results have recently been developed in the discrete-time framework [46], [87].

